# Factors associated with adherence to annual rabies vaccination in dogs and cats in the municipality of Curuçá, Eastern Amazon

**DOI:** 10.1101/2022.11.15.516558

**Authors:** Elane A. Andrade, Kelly K. G. Nascimento, Mateus B. Silva, João V. Morais, Mario J. Carneiro, Maiara V. Monteiro, Carolina F. Azevedo, Christiane M. B.M. Rocha, Luciana B. Chaves, Karin C. Scheffer, Rene S. Cunha Neto, Isis Abel

## Abstract

Dogs and cats maintain the urban cycle of rabies, and vaccination is the main form of prevention and control of the disease. Brazil has seen human rabies cases transmitted by dogs and cats infected with the bat variant in regions where annual campaigns no longer take place. Although the municipality of Curuçá has no cases of urban rabies and viral circulation in wild animals is unknown, there are informal reports of contact of animals and people with vampire bats. This study aimed to identify factors associated with immune response against the rabies virus in dogs and cats in this municipality. A total of 352 dogs and 46 cats were randomly selected for blood collection and their owners answered a questionnaire. The animals were mostly males, aged between 1-3 years, and with unrestricted access to the street. A total of 48.8% of dogs and 32% of cats were not vaccinated in the last anti-rabies campaign, and 4.7% of dogs had been attacked by bats. Among the analyzed samples, only 21.1% had a titration ≥ 0.5 IU/mL. Risk factors for not participating in vaccination campaigns included species, presence of veterinary care, and participation in annual vaccination campaigns (OR = 0.46, 2.55, and 15.67 respectively). The animal population was estimated at 18,620 dogs and 4,556 cats. The human:dog ratio was 2.1:1 and the human:cat ratio was 8.7:1. This study revealed that the estimated population of dogs based on the human population was an underestimate for communities in the Amazon region. This was the first time that the number of dogs attacked by bats was determined. Health education with an emphasis on responsible ownership and periodic and biannual rabies vaccinations are recommended for the municipality.

**AUTHOR SUMMARY:** Rabies is a viral disease characterized by brain and spinal cord inflammation. It affects all mammals, being almost 100% lethal. Hematophagous bats are one of the main wild reservoirs responsible for outbreaks of human rabies in the state of Pará and other regions of Brazil. Vaccination is the most effective form of control and prevention, even where rabies is believed to be under control. The levels of antibodies that fight the rabies virus must be constantly monitored through serological analysis to assess the effectiveness of vaccination programs. In the municipality of Curuçá, Pará, bats commonly attack people and their pets, and anti-rabies campaigns are not promoted every year. In the present study, we determined the number of attacks by bats on domestic dogs and estimated vaccination coverage and the canine and feline population in the municipality, which was underestimated. This information can be useful for future vaccination campaigns. The study identified factors associated with responsible ownership that interfere with the protective titration of animals against
 rabies and suggests promoting more than one annual anti-rabies campaign in this location.

## INTRODUCTION

Stray or semi-domesticated dogs and cats are the main sustainers of the urban cycle of rabies, and when not vaccinated, they become important sources of infection for humans ^[1]^.

Despite the decrease in cases of canine rabies since 1970 in Brazil ^[2]^, between 2000 and 2009 approximately 4,177,409 human rabies consultations were reported in the country. Data from the Notifiable Diseases Information System (*Sistema de Informação de Agravos de Notificação* - SINAN) show that among animals, dogs were the main human aggressor ^[3,4]^.

Between 2007 and 2017 this condition increased in several regions of the country, ^[5]^ including the state of Pará. In the micro-region of Salgado, northeastern Pará, 74.1% of the notifications in SINAN came from aggression by dogs ^[6]^.

The most recent cases of human rabies in Brazil were caused by the AgV3 variant, which is typical of bats, but is transmitted by cats, demonstrating that there is contact between bats and domestic animals and between bats and humans ^[7]^.

In the municipality of Curuçá, there are no records of human or canine rabies and the circulation of the rabies virus has not yet been studied. However, the lack of records does not necessarily indicate the absence of viral circulation, but rather that the virus may be present in silent areas ^[8,9]^.

It is known that in this municipality, in 2012 and 2013, about 10% of rabies treatment was required following attacks by bats against people living in rural areas, where domestic animals walk roam freely, and are under favorable conditions for this type of contact. ^[10,6]^.

Considering the information above, this work aimed to investigate the epidemiological factors associated with the presence of neutralizing antibodies against the rabies virus in dogs and cats, vaccinated or not, owned in the municipality of Curuçá-Pará.

## METHODOLOGY

### Ethical aspects

The research project was approved by the Ethics Committee on Animal Use (*Comissão de Ética na Utilização de Animais* - CEUA/UFPA), under No. 6363180918.

### Study area

The municipality of Curuçá is located in the northwest of the Brazilian Amazon region, 130 km from Belém. More than half of its territory is part of the Mãe Grande de Curuçá marine extractive reserve, which consists of an environmental conservation unit that was established to ensure the sustainable use of its resources, which are used by most of its inhabitants for subsistence. This municipality includes the Curuçá River and a large coastal area mixed with Amazonian riverside zones, mainly occupied by mangroves. Due to its location, the municipality’s economy is based on fishing, and trade of fish and shellfish collected in the environment.

### Sample size

This research was based on the estimate of the Ministry of Health (*Ministério da Saúde* – MS; presented in the resolution No. 05/2013) that the canine population corresponds to 20% of the human population. Therefore there are approximately 6,859 dogs in Curuçá, based on the last human census (34,294 inhabitants - IBGE, 2010). The required canine sample size was calculated using the Statcalc tool of the Epi Info™ software version 7.2, with an expected frequency of 60% and a significance level of 5%, which resulted in a total of 350 animals.

### Sampling method

The dogs were selected by random sampling, prioritizing the evaluation of one animal per household. The households were selected using the QGIS 3.2.2 software. The selected points were located using the MapIt-GIS Data Collector application. If there were no dogs in the randomly selected household or it was empty, the next household was promptly selected (Figure 1). Cats were selected for convenience in randomly selected households where dogs and cats were present.

**Figure 1.**
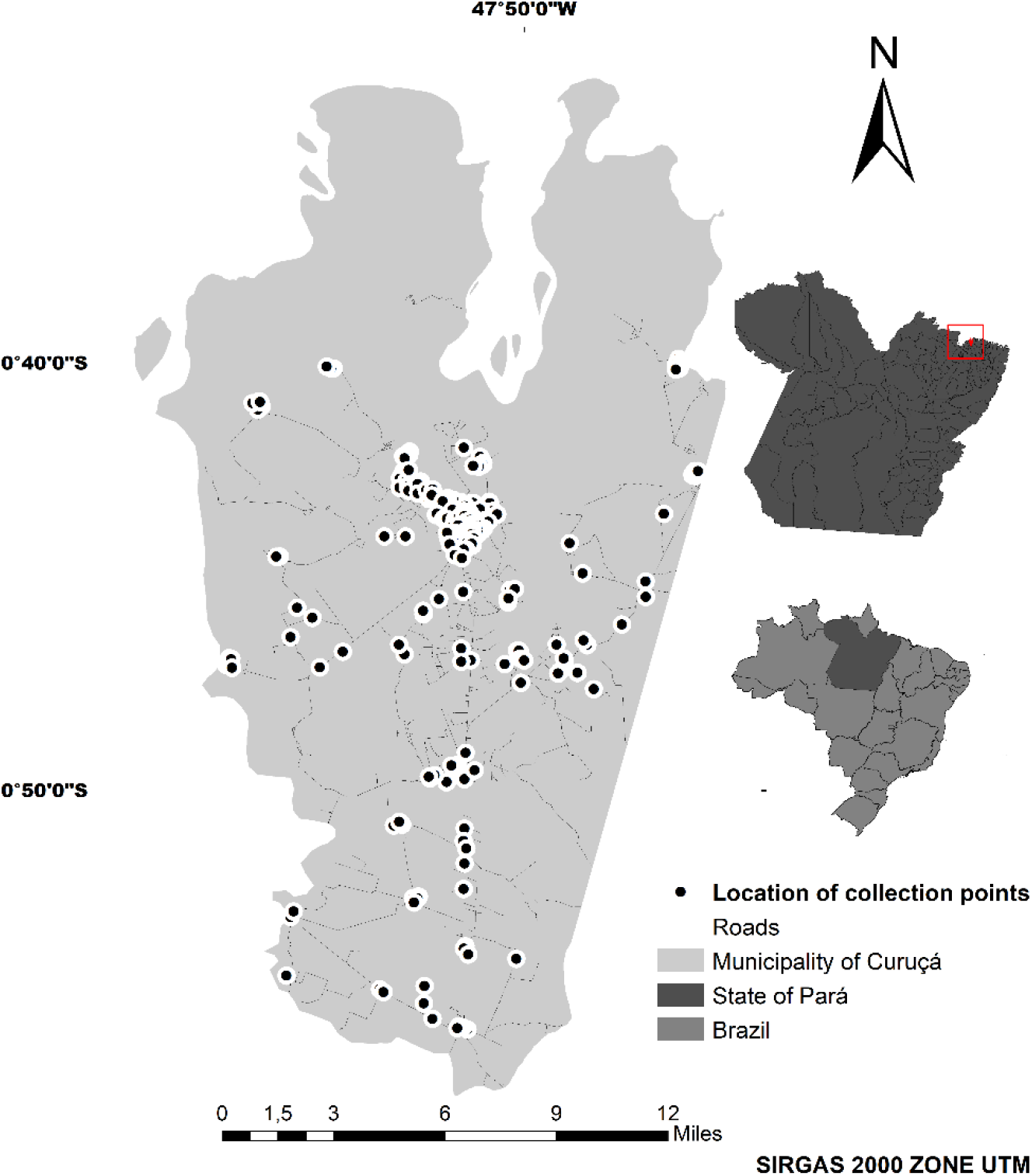
Location of the study area and distribution of random points for collecting biological material from dogs.

### Questionnaire administration

After signing an Informed Consent Form and adhering to the project protocol, owners were invited to answer a semi-structured questionnaire containing items identifying the animal, daily habits, and general care.

### Collection of biological material

Whole blood was collected by a professional veterinary using a closed collection system or a needle-punched syringe, through puncture of the cephalic or external jugular vein, in a volume of 3 to 5 ml. Samples were placed in plastic tubes for blood collection and subsequent serum collection. Serum samples were stored in a freezer at −20°C until use.

### Serological analysis

Samples of 374 animals were sent for serological analysis. Four canine samples arrived in poor condition and could not be evaluated. Thus, 370 samples were analyzed; 324 from dogs and 46 from cats.

The Rapid Fluorescent Focus Inhibition Test (RFFIT) was performed for measuring neutralizing antibodies against the rabies virus ^[11]^, with modifications.

First, the received sera were inactivated in a water bath at 56°C for 30 minutes and then six serial dilutions of each serum were carried out, in the ratio 2, starting from 1:5. A total of 25 μL of serum was placed in the first opening with 37.5 μL of Eagle’s Minimum Essential Medium (MEM) with Earle’s salts, and it was supplemented with 10% inactivated fetal bovine serum.

After dilution, 50 μL of virus of the Challenge Virus Standard (CVS) strain were added, and the plates were incubated in an oven with CO2 at 37°C. After 1 h 30 min of incubation, 50 μL of BHK-21 cells (2.5 × 104 cells/mL) were added and incubated again at 37°C in an atmosphere containing 5% CO_2_ for 20 hours. The cells were fixed in an ice bath and the reaction was revealed by adding anti-rabies virus conjugate. The reading was performed under a LEICA^®^ DMIL inverted fluorescence microscope at 200 X magnification. The titers were calculated using the Spearman-Kärber method of analysis, by comparison with the standard serum, and the results were expressed in International Units per milliliter (IU/ml), with animals with results equal to or greater than 0.5 IU/ml being considered immune and those with lower titers being considered non-immune.

### Data analysis

#### Statistical analysis

Descriptive analysis of the variables addressed in the companion animals’ questionnaire was performed using the SPSS v.24 software. The chi-square test (χ^2^) was used to test the associations between the variables. Associations were significant when p<0.05. In these cases, the odds ratio (OR) was calculated.

Variables with significant associations and those with P ≤ 0.02 were included in the binary logistic regression model, using the “ENTER” method, with the following dependent variables: “was vaccinated last year”, “usually vaccinates annually” and “result of serology”. All covariates raised in the questionnaire were tested.

#### Calculation of population estimate

Sampling fraction methodology was used to estimate the population of dogs and cats in the municipality of Curuçá ^[12]^, with modifications; where the ratio between the total number of animals counted in the visited households and the sample fraction was estimated, according to following formula:

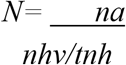

Where: *N*= number of estimated animals in the municipality; *na* = number of animals counted in the visited households; *nhv* = number of households visited; *tnh* = total number of households in the municipality.

## RESULTS

### Characterization of the animal population

A total of 352 dogs were selected. Of them, 56.7% were males and 43.3% were females, with most of them being aged between one and three years (48%) and being of mixed breed (93.8%).

Although most were companion animals (68.4%), there was also a percentage of guard animals (24.6%) and some intended for hunting (5.7%) and fishing (1.3%). The majority animals were adopted (76.4%), followed by those born at home (17.8%). As reported by the owners, 81.8% of the animals had access to the street on their own, spending the day outside the home and at night they returned and slept within the property (83.1%), either on the balcony (42.5%), in the yard/ open sky (45.5%) or in a dog house (12%). Also, among those who had free access to the street, 16.9% slept inside the house.

Most owners (68.5%) reported that they usually take their animal to the rabies vaccination campaign every year; however, 48.8% of the animals had not been vaccinated in the previous year. Among the reasons why not, some owners reported that the animal was too young to receive the vaccine or had not yet been born (44.4%), and others reported other reasons (Figure 2).

**Figure 2.**
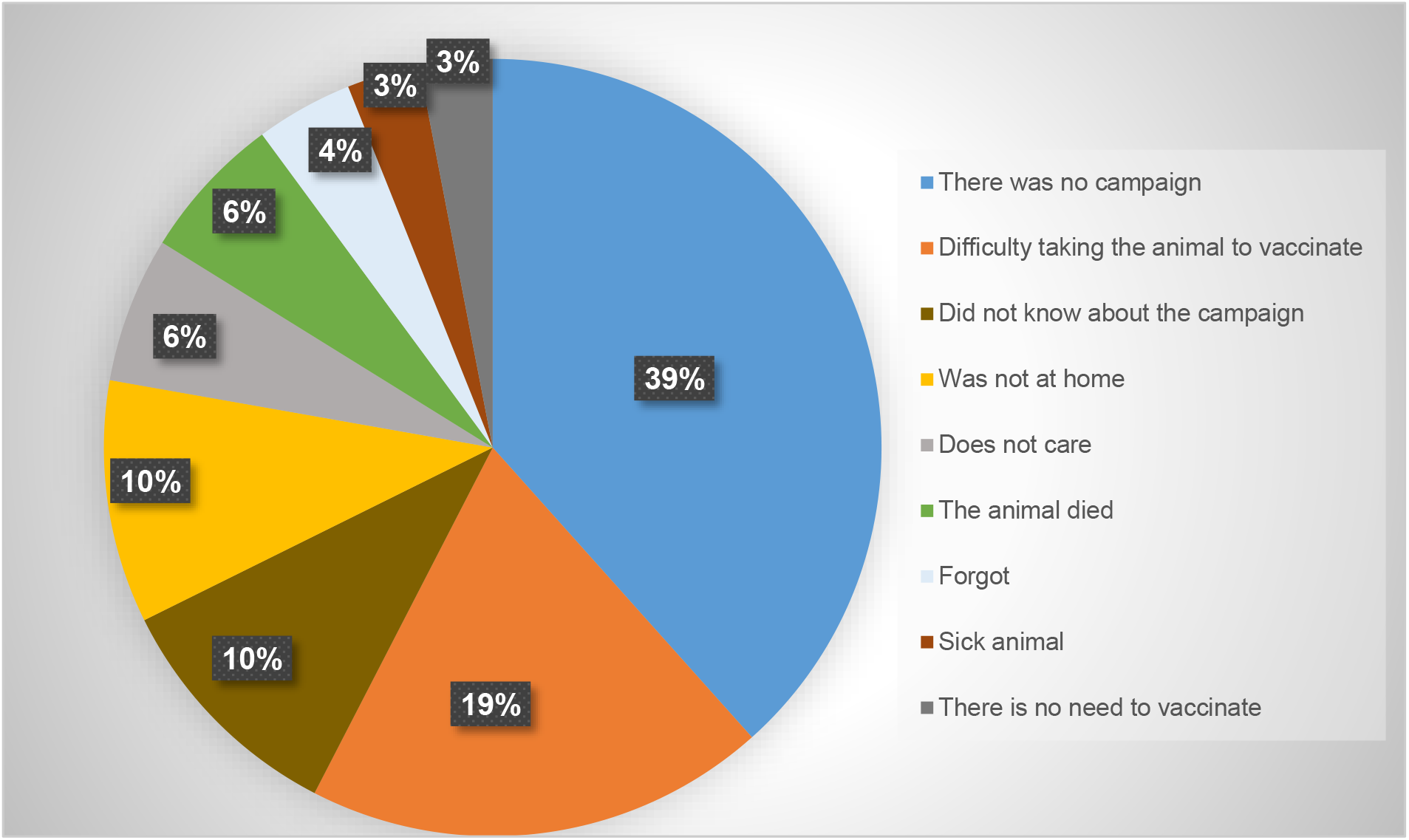
Reported reasons for why dogs were not vaccinated in the previous year.

A total of 4.7% of dog owners reported that their dogs had been attacked by bats at least once in their lives. These aggressions mostly occurred on islands in the municipality, where dogs accompanied their owners on fishing activities.

In the present study, 46 cats were selected, most of them being male (69.6%), aged from one to two years (44.4%), adopted (89.1%), having access to the street (89.1%), but sleeping inside the house (54.3%). About 32% of the owners reported not having vaccinated their cats in the last anti-rabies campaign that took place in the municipality. When asked about the reasons why not, 53.4% reported that they still did not have cats, or that their cat was too young to receive the vaccine. The others reported other reasons, which are described in Figure 3.

**Figure 3.**
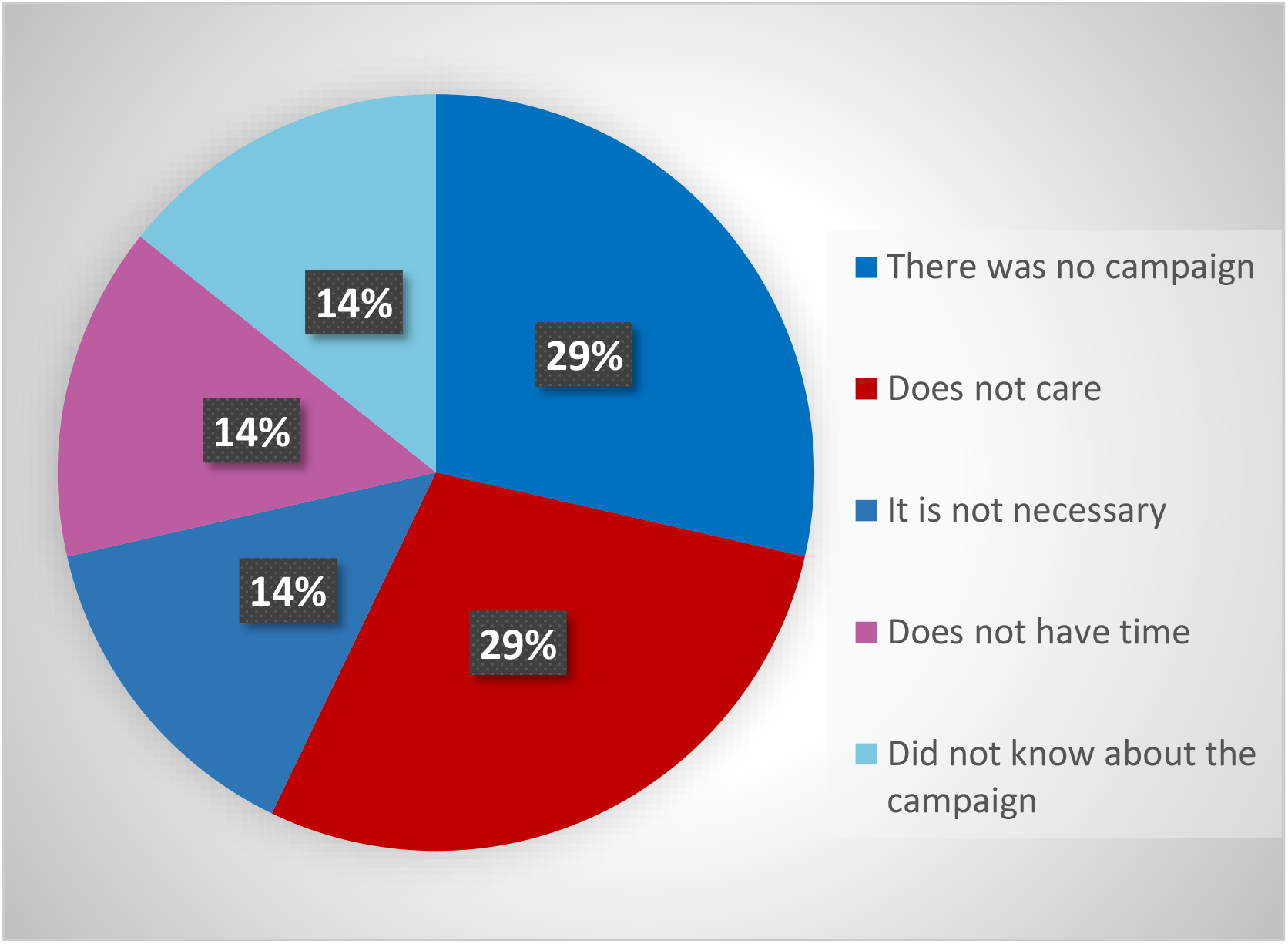
Reported reasons for why cats were not vaccinated in the previous year.

Animals that did not receive veterinary care such as deworming and other vaccines and whose owner was not used to adhere to anti-rabies vaccination campaigns in the municipality were more likely to have not been vaccinated in the previous year. However, being a dog (OR=0.5) and having another role besides companionship (OR=0.6) were protective factors for these animals to be vaccinated in the previous year. The study also revealed that the older the animal, the greater the chance of being vaccinated (Table 1).

**Table 1.**
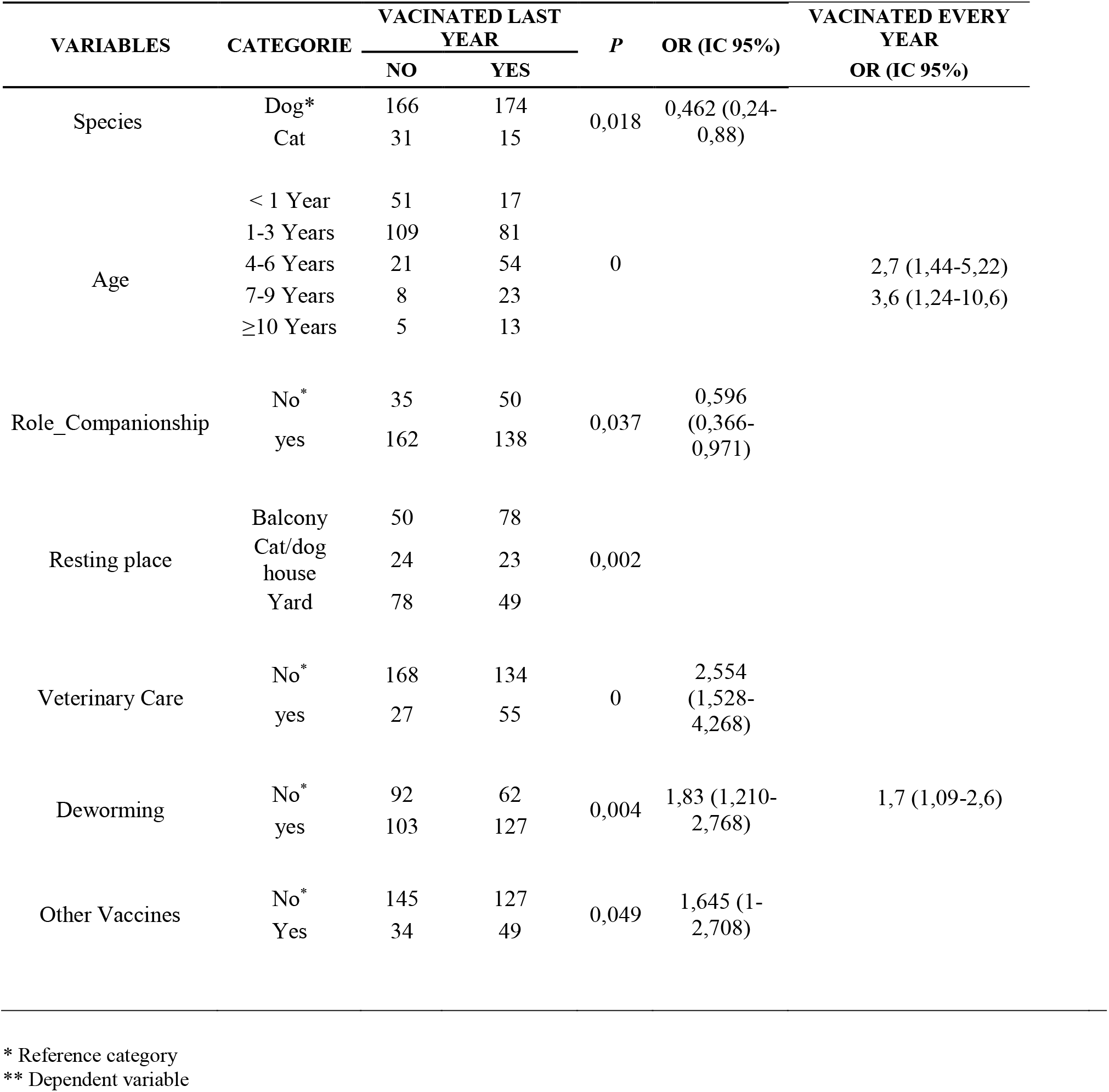
Factors associated with owners’ reports of rabies vaccination of their dogs and cats in the last campaign in the municipality of Curuçá, Pará.

When evaluating the candidate prediction variables, with rigorous annual vaccination as the dependent variable, the analysis revealed that older animals that do not receive care other than food are more likely to not participate in vaccination campaigns.

### Serological analysis

Of the total samples analyzed, only 21.1% showed protective titration.

Non-protective titration was present in 63.9% and 69.6% of the dog and cat samples, respectively.

When considering only dogs and cats that usually received the rabies vaccine every year in the campaign, only 61 (26.5%) dogs and 8 (33.3%) cats had titers ≥ 0.5 IU/ml. Moreover, seven unvaccinated dogs and two cats had positive titers. As for the animals that received the vaccine in the previous year, most (58.7%) did not have sufficient protective titration.

The analyses revealed that age was a risk factor for the absence of rabies titration, with increasing age being associated with a greater chance of having a negative titer (< 1 year: OR=0.43 CI 95%=0.006-0.320; 7-9 years: OR=8.66 CI 95%=3.59-20.88). Other associated risk factors were the place where the animal rests, lack of veterinary care, not participating periodically in anti-rabies campaigns, and non-vaccination in the last campaign, the latter having about seven times the chance of not being protected (Table 2).

**Table 2.**
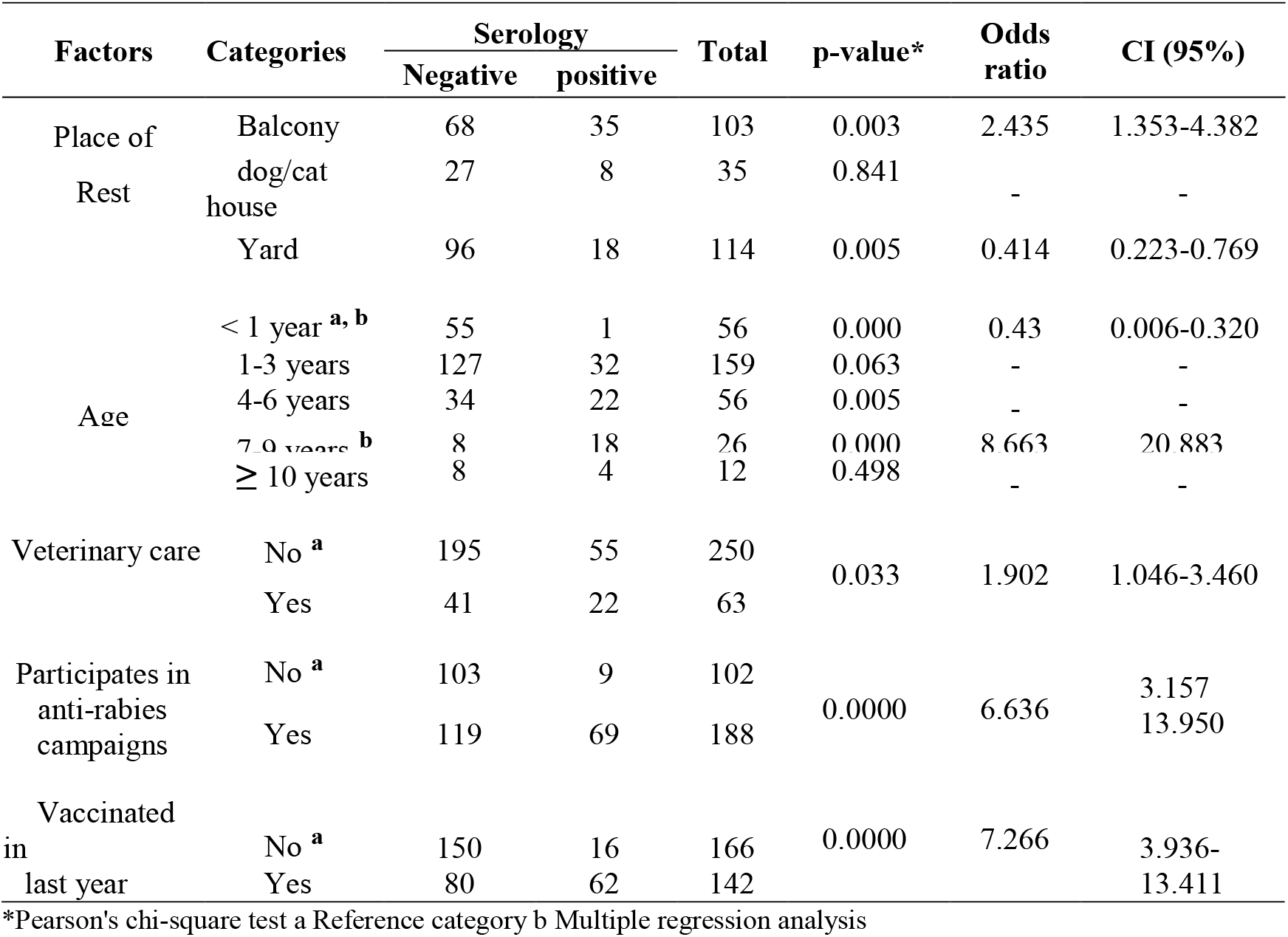
Analysis of associations between the variables of the questionnaire and the result of the rabies serological teste.

### Estimation of the population of dogs and cats in households

Considering the numbers of dogs (703), cats (172), and households (321) selected for this study and the number of households in the municipality according to the last IBGE census (8,502), the animal population was estimated at 18,620 dogs and 4,556 cats, with an average of 2.19 and 0.53 dogs and cats per household, respectively.

In 2019, the human:dog ratio was 2.1:1 and the human:cat ratio was 8.7:1, that is, in the municipality of Curuçá, there was one dog for every two people and one cat for every eight people.

## DISCUSSION

Dogs and cats are the most prevalent domesticated animals in the world, and they coexist in great intensity with the human population ^[13]^. This interaction can cause the exchange of pathogens, leading to the development of zoonoses, including rabies, in human populations ^[14]^. It is estimated that the global load of dogs in the world reaches 700 million animals ^[15]^. Brazil is the second largest country with respect to its number of domestic animals, with approximately 52.2 million dogs and 22.1 million cats ^[16]^.

Most of the animals studied in both species were male, as reported in other studies ^[17,18,19,20]^. This suggests that the human population has a preference for this gender since females can become pregnant which can increase the number of animals, and the methods of sterilization of females are more expensive ^[20]^.

Despite the large percentages of dogs and cats for companionship (68.4% and 100%, respectively), these animals are considered free-living or “from the neighborhood”; that is, they are semi-dependent and semi-restricted or even unrestricted. This means that although they do not always depend directly on the human to provide food and shelter, they have a reference family ^[21]^.

The animal population is considered young, between 1 and 3 years old (48%), corroborating a study by Magnabosco ^[22]^ in São Paulo. Most (84%) had free access to the street, as reported in other studies in Brazil and other countries ^[17,18,23]^, and which can increase the risk of accidents. Therefore, the animal population has a shorter lifespan and will constantly renew as new animals arrive. The free access to the street also allows contact with wild animals, increasing the chance of aggression and transmission of pathogens to humans ^[24]^.

Studies show that most animal bites are caused by dogs and cats ^[25,6,26]^ from the neighborhood or even belonging to the victim ^[27,28,26]^, in unprovoked conditions. In the present study, about 22% of dogs and cats, despite spending the day outside the house, sleep together with the family, inside the house, which increases the chances of aggression and transmission of pathogens to humans ^[23]^. In this context, controlling animal movement is extremely important to reduce the risk of exposure to rabies.

In Curuçá, these domestic animals are also found in wild territory, since 22 (5.7%) dogs were used by their owners to assist in hunting. This practice is common in many cultures in the world ^[29,30]^, and involves letting the animal run free in search of prey ^[31]^, without constant supervision by the hunter. At this moment, there is a connection between domestic and wildlife, allowing the transmission of pathogens between species ^[32,15]^ and involving the human population in this connection ^[33]^. Moreover, interaction between hunters from different places and their dogs enables the transmission of pathogens between communities ^[31]^.

Given its location and economic base, some residents of extractive reserve (RESEX) develop fishing activities and many take their dogs, which explains the finding that the main role given for five animals was fishing. In this study, about 4.7% of the dogs had already been attacked by bats. Aggression occurs when people together with their animals move long distances, using temporary makeshift shelters and staying several days without protection against bats ^[34]^, or when the animals live together with their owners in areas of mangrove forests ^[35]^.

The study by Hughes & Macdonald ^[15]^ showed that, in several parts of the world, domestic dogs constantly interact with several wild species, including bats, either for predation, which can be in either direction, or in the transmission of pathogens.

Cats are also important in this context since most of them have access to the street (89.1%) and sleep inside the house (54.3%), in addition to being hunters by nature. According to Welch & Leppanen ^[36]^ cats are the species most responsible for attacks on bats, whether for predation or for fun ^[37]^.

Cats come into contact with bats in their shelters ^[38]^, or when they come across animals that fall while flying during the day ^[22]^. According to Castilho et al. ^[39]^ this type of contact is responsible for cases of rabies in this species.

It is worth mentioning that the most recent cases of human rabies reported in Brazil had cats as transmitters, with one case in Jacaraú-PB, in 2015 ^[40]^, and two more cases in Boa Vista-RR (2016) and Recife-PE (2017) in the following two years. In all these cases, the variant involved was AgV3, common in vampire bats. In 2019, there was a case in Santa Catarina, with transmission by a domestic cat, and in 2020 a case in Paraíba involving the AgV2 variant transmitted by a wild canid. Thus, there are predisposing conditions for the occurrence of human cases in the studied municipality.

In addition to establishing pre-exposure immunity, rabies vaccination of domestic animals protects individuals, and thus can prevent the spread of the virus to humans and other animals ^[41]^. The state of Pará, among other states, has canine vaccination coverage below 70% ^[16]^. Similarly, according to the present study, the municipality of Curuçá vaccinated 51.2% and 32.6% of dogs and cats (respectively) in the most recent rabies vaccination campaign, conducted in 2018.

These results are below the level recommended by WHO and the levels described in other regions of the country ^[22,42,16]^ and are above levels reported in countries where urban rabies is considered endemic. In locations such as the Philippines, Mali, and the Democratic Republic of Congo, vaccination coverages ranged from 17% to 47% ^[20,43,44]^.

The reasons for low vaccine rates most frequently reported in other locations were coverage, negligence, the belief that the vaccine is not effective, lack of time, lack of knowledge, distance from vaccination posts, and fear of adverse effects from the administered vaccine ^[45,20,46]^.

In Curuçá, the rabies vaccination campaign takes place on a “D” day, where fixed posts are set up in the central region of the city and receive dogs and cats for vaccination. On other days, vaccination takes place door-to-door. However, the campaign is conducted in a decentralized manner, with each health unit being free to conduct the vaccination campaign in the way it deems most appropriate for its area of coverage. In some locations, this does not occur at all due to difficult access, so dog owners reported that vaccination did not take place in that area (39%) or that due to the distance there was some difficulty in taking the animal to the vaccination post (19%). Besides this situation, another 10% reported not having been informed about the vaccination campaign that year, reinforcing that communication efforts must be made constantly in these populations ^[47]^.

Some cat owners also reported no campaign in their location (29%). However, what stands out is the fact that they do not care about vaccinating their animals (29%) and believe that the species does not need to be vaccinated against rabies.

In the study by Kongkaewa et al. ^[17]^ a part of the population reported that vaccination was useless in cats, as they did not believe that the species was also prone to rabies infection.

The absence of canine and feline rabies cases in the municipality may also contribute to the low adherence to vaccination, as according to Rodrigues et al. ^[42]^ this absence generates a false sense of protection in the population and health agencies. The municipality of Curuçá conducts rabies vaccination campaigns annually, around the months of August and September. However, in 2016 and 2017 there were no campaigns according to the Information System of the National Immunization Program (*Sistema de Informações do Programa Nacional de Imunizações* - SI-PNI) ^[48]^, which may have discouraged the population from vaccinating their animals.

About 6% of respondents reported not taking their pets to be vaccinated because, in their perception, when vaccinated against rabies, dogs and cats get sick and die. In fact, in 2010, the anti-rabies campaign was interrupted after successive deaths of animals with neurological signs. Until that year, the vaccine used in the campaigns was Fuenzalida & Palácios, consisting of an inactivated virus produced in the nerve cells of mice ^[49]^. It was suspected that batches of the vaccine were compromised, as it has been previously associated with Guillain-Barré syndrome in humans ^[50]^.

From 2010, the Ministry of Health changed the vaccines distributed in the dog and cat campaigns to a cell culture vaccine, which provides greater safety and efficacy in the conversion of protective titers in these animals ^[2]^.

Possibly because of what happened, the population has become more resistant to the administration of vaccines to their animals and many have even reported this fact, which may also be associated with the high chance of dogs not having been vaccinated in the anti-rabies campaign in the previous years.

Moreover, participation in the vaccination campaign was closely associated with the interaction between the animals and their owners, since youngest animals were the ones more likely to be vaccinated, as reported by Mauti et al. ^[46]^. Culturally, in this region, puppies are likely to receive more attention from their owners because at this stage they are more closely associated with the children in the house. This information is aligned with the association revealed by the study, that companion animals and those who receive other care such as veterinary care and periodic vaccination were more likely to have been vaccinated in the previous year, corroborating what was reported by Davlin et al. ^[20]^, that the chance of vaccination was greater among domiciled animals, due to the constant proximity between the animals and human beings.

The seroprevalence of antibodies against the rabies virus is quite heterogeneous, as it depends on the method of analysis, the animals involved, and the region where these animals are located ^[51,52,53]^. In the present study, the global seroprevalence of neutralizing antibodies was around 20%, which, despite being low, is in accordance with other reports for farm animals ^[46]^. Although there are reports that cats have higher titers than dogs, possibly due to particularities of the immune system that are not yet well understood ^[53]^, the present study did not show any differences in immune response between species (p=0.97).

However, it was possible to notice a difference between vaccinated and unvaccinated animals in the previous year, with most dogs and cats that participated in the vaccination campaign not having a protective titer and some unvaccinated animals having neutralizing antibody titers ≥0.5 IU/ml.

The low titer of neutralizing antibodies in vaccinated animals may be associated with factors such as the number of doses received by the animal in its lifetime ^[54,55,56,57]^, the time between vaccination and collection for analysis ^[53,58,59]^, the age of the animal that received the vaccine ^[60]^, and the type of vaccine used ^[61,41]^, because according to Kennedy et al. ^[62]^, the differences in the formulation and production of different vaccines can produce different failure rates and immune responses.

The time elapsed between vaccination and collection for serological testing can also influence the response to antibody production, as the chances of testing positive tend to decrease as this period increases. The ideal time to perform the collection is between 4 and 8 weeks after vaccination ^[53,58,59]^, and the collections of the present study took place between 10 and 12 months after the last rabies vaccination campaign, and therefore may be associated with the small proportion of animals with positive titer.

According to Gold et al. ^[63]^, multiple exposures to small doses of the virus can stimulate the development of antibodies, preventing future infections. This type of non-lethal exposure has been reported in humans in an isolated indigenous tribe who never received rabies prophylaxis and were exposed to bat attacks ^[64]^. People and their animals in Curuçá are in constant contact with hematophagous bats, which can make the local population acquire antibody titers, and helps to explain the 7.6% animals who have never been vaccinated and yet produced antibodies at protective levels.

Moreover, the sensitivity and specificity of antibody detection methods can impair the detection of protective titers ^[63]^ and the presence of antibodies can result from an alternative course of exposure, where the elimination of the virus by the organism occurs even before it invades the CNS, causing a subclinical infection but with antibody production ^[65,66]^.

Young and old animals are more prone to immunological failures, as the former do not have an immune system mature enough to produce an adequate response and the latter are already entering the cellular senescence phase, decreasing their immune capacity ^[41,67]^.

Some of the animals in the study were not vaccinated because they were newborn, and one year later it would be expected that they would not have a titration. As older animals are less likely to be vaccinated every year, it is possible that many of them received only one dose as puppies, which could explain the fact that age was a risk factor for the absence of rabies titration, and younger and older animals had a greater chance of not showing protective titration.

It is recommended that puppies receive a booster dose after primary vaccination and periodic reinforcements from then on because even with these animals being born from vaccinated bitches, they cannot acquire sufficient titration for protection even 30 days after vaccination ^[55,57]^.

This study showed that the animals in Curuçá received only the essential for their survival, such as food, water, and shelter, but not veterinary care nor other vaccines. Although the rabies vaccine is available throughout the year in agricultural stores, pet shops, and food stores, most of the population only seeks the vaccines donated through campaigns, which take place once a year throughout Brazil.

When planning a vaccination campaign, counting or estimating the population of animals is essential, as it allows the identification of the necessary resources and the best methods ^[68]^. Currently, in most of Brazil and other countries, the estimation of animal population for rabies vaccination campaigns is based on the calculation of the human population ^[69]^. However, adopting a single ratio or percentage for population estimates for a continent can lead to errors in future, as each location has particularities that must be taken into account ^[23,16]^. This study presents population estimates and epidemiological characterization of dogs and cats in a municipality in the state of Pará and evaluates the titration of antibodies against rabies in animals, which can help vaccination and disease control programs in the municipality and the state.

According to data provided by the 3rd Regional Health Center of the Public Health Department of the State of Pará (*3° Centro Regional de Saúde da Secretaria de Saúde Pública do Estado do Pará* - 3°CRS/SESPA), in 2018 the municipality of Curuçá reached 136.3% and 97.5% of the vaccination goal for dogs and cats, respectively. However, such results close to and above 100% are common and were obtained because the anti-rabies vaccination campaigns in Brazil are programmed and organized while considering the information established by WHO ^[21]^ and the Pasteur Institute ^[49]^, which estimate the human:dog ratio at 10:1 and 7:1, and also by the Ministry of Health, which estimates that the canine population corresponds to between 10 and 20% of the human population.

Data from the present study revealed that the human:dog ratio in the municipality of Curuçá was 2:1, that is, the canine population corresponded to 50% of the human population. These results are far from the recommended for emerging countries by the WHO, of one dog for seven or eight inhabitants ^[21]^. In studies conducted in Southeastern Brazil, this relationship varied by location. Studies have reported the human:dog ratio as 7:1 ^[70]^, 5:1 ^[71,72]^, and 4:1 ^[73,18]^. These ratios can be considered while planning vaccination campaigns in the corresponding regions, but should not be used for planning vaccination campaigns in other regions of the country. If the numbers of the present study are taken into account, together with the fact that only 51.2% of the studied dog population was vaccinated in the previous year, it can be inferred that the 80% vaccination coverage recommended by the WHO was not achieved, as also described by De Lucca et al. ^[74]^ and Rodrigues et al. ^[42]^ in São Paulo, and Fernandes et al. ^[75]^ in Santa Maria/Rio Grande do Sul.

Until 2018, due to delays in the delivery of vaccines and in the programming of campaigns, the vaccination coverage rates varied greatly in almost all Brazilian municipalities. In 2019, the campaign took place only in higher risk areas, in the states of Maranhão, Ceará, Rio Grande do Norte, Mato Grosso do Sul, Mato Grosso, Rondônia, and Acre ^[2]^.

Southern Brazil is considered controlled for urban rabies ^[16,75]^, which led to the cancellation of anti-rabies campaigns in the region since 1995. Until 2015, only the state of Paraná promoted campaigns and only in municipalities bordering Paraguay ^[2]^.

However, the absence of vaccination has caused animals to lose the immune reinforcement provided by it and, despite the large decrease in cases of canine and feline rabies since 1999, there was an increase in the infection of dogs and cats ^[75]^ with consequent transmission to humans. This demonstrates the importance of these animals as secondary transmitters and of maintaining vaccination even where the disease has already been eliminated ^[76]^. Furthermore, the duration and size of outbreaks tend to be longer in unvaccinated populations ^[77]^. It is worth mentioning that the most recent cases of human rabies were reported in Santa Catarina, where it was transmitted by domestic cats in 2019, Angra dos Reis/Rio de Janeiro, where it was transmitted by bats, and Catolé do Rocha/Paraíba, where it was transmitted by domestic dogs, both in 2020 ^[2]^.

Therefore, the maintenance of current campaigns is suggested, in order to achieve both a greater proportion of truly immunized animals ^[62]^ and the promotion of education programs for responsible ownership, since some factors associated with general care are associated with the acquisition of antibodies against the rabies virus.

## CONCLUSION

This study revealed that the estimate of the population of dogs based on the human population was an underestimation for communities in the Amazon region, where the animals were semi-restricted or unrestricted, having free access to the wild environment. This was the first time that the number of dogs attacked by bats was determined. These dogs are indicators of the possible re-emergence of urban rabies in the region. Factors associated with responsible ownership were identified as risk factors for periodic vaccination of these animals and, therefore, the region should promote health education with an emphasis on responsible ownership and periodic and biannual rabies vaccination.

## ACKNOWLEDGEMENTS

The authors thank Capes, Cnpq and PROPESP/UFPA (PAPQ) for their financial support and the Pasteur Institute for the serological analyses.

